# Revalidation of *Manis aurita* based on integrative genomic and morphological evidence

**DOI:** 10.1101/2025.07.05.663294

**Authors:** Narayan Prasad Koju, Zeling Zeng, Guihua Zhang, Zhicheng Yao, Xia Huang, Xiaoyun Wang, Melissa T.R. Hawkins, Arlo Hinckley, Mary Faith C. Flores, Ce Guo, Jun Li, Devendra Maharjan, Saraswoti Byanjankar, Lianghua Huang, Wenhua Yu, Liang Leng, Kai He, Anderson Feijó, Yan Hua

## Abstract

Pangolins face severe conservation threats globally, necessitating accurate taxonomy for effective conservation. Previous genomic research on the Chinese pangolin (*Manis pentadactyla*) identified two deeply divergent lineages (MPA and MPB), suggesting underestimated species diversity. The recent description of *M. indoburmanica* (corresponding to MPB), however, did not assess its relationship with historical names, particularly the subspecies *M. p. aurita*, leaving the group’s taxonomic status uncertain. To resolve this issue, we employed an integrative framework, analyzing genomic and morphological data from museum specimens including the lectotype of *M. p. aurita* to clarify phylogenetic relationships and taxonomy within *M. pentadactyla* sensu lato. Our results demonstrate that MPB, which includes both *M. indoburmanica* and *M. p. aurita*, is geographically restricted to the southern Himalayas and thus distinct from other *M. pentadactyla* populations. Genomic analyses indicate the two clades diverged ∼1.8 Ma and have remained largely isolated, with only minimal gene flow with congeners. Furthermore, morphometric analyses of both cranial and external features reveal consistent and significant differentiation between the Himalayan lineage (MPB) and *M. pentadactyla* sensu stricto. Collectively, these congruent findings provide unequivocal support for the revalidation of *Manis aurita* Hodgson, 1836, thereby establishing it as a distinct extant species of Asian pangolin.

## INTRODUCTION

Pangolins (Order Pholidota) are unique mammals characterized by their keratinized scales, an extraordinary trait among mammals ^1^. These animals face severe threats including habitat loss and extensive illegal trade, which has resulted in dramatic population declines ^2–4^. Consequently, all pangolin species are listed under Appendix I of the Convention on International Trade in Endangered Species of Wild Fauna and Flora (CITES) and are recognized as Evolutionarily Distinct and Globally Endangered (EDGE) species ^5^. Accurate species identification, clarification of species distribution, and estimation of population sizes and genetic load are essential for effective conservation efforts ^6–8^.

Until recently, eight extant pangolin species were recognized: four in Africa and four in Asia ^9^. All Asian species belong to the genus *Manis*, including Chinese pangolin (*Manis pentadactyla*), Indian pangolin (*M. crassicaudata*), Sunda pangolin (*M. javanica*), and Philippine pangolin (*M. culionensis*). Recent genomic studies revealed cryptic genetic and taxonomic diversity within Asian pangolins. For instance, genome sequencing of confiscated scales uncovered a cryptic lineage, provisionally named “mysteria”, for which no voucher specimens or precise locality data exist ^10^. Moreover, analyses of broad sampling within the Sunda pangolin ^11,12^ and *M. pentadactyla* ^10,13,14^ have revealed distinct clades, indicating the existence of an undescribed species.

The Chinese pangolin is distributed across a broad range, extending from Nepal in the west to the eastern coast of China, with insular populations on Hainan and Taiwan islands. Latitudinally, its range extends from central China southward to Southeast Asia ^15^. This broad distribution is divided by geographic barriers, such as the mountains of southwestern China and southward-flowing river systems, which likely drive genetic divergence among geographic populations ^16,17^. Accordingly, Hu, et al. ^13^ identified two major clades within *M. pentadactyla,* namely, MPA and MPB. While recent studies have found additional subclades closely related to MPA in China and Southeast Asia ^14,18^, and MPB remains the most genetically divergent lineage, and its geographic origin has been enigmatic. Initially, MPB was known almost exclusively from specimens confiscated along the Sino-Myanmar border, suggesting a Myanmar origin. This inference was tentative, as the area is a major wildlife trafficking hub ^19–23^. However, recent studies found occurrences of MPB in Nepal, South Tibet and northeastern India ^24–27^, strongly suggests that this clade represents the western populations inhabiting the South Himalayan region.

Beyond its geographic origin, the formal nomenclature and taxonomic identity of the MPB clade remain unresolved issues. Historically, three subspecies of *M. pentadactyla* were recognized: the nominate subspecies *M. p. pentadactyla* Linnaeus, 1758 described based on specimens from Taiwan; *M. p. pusilla* Allen, 1906 from the Hainan Island; and *M. p. aurita* Hodgson, 1836 from the central Nepal ^28^. While the former two have been subject to extensive genetic studies ^13,14,18,29,30^, and their evolutionary relationships as well as distribution across China and SE Asia have been confirmed ^13,14,31^, the taxonomic status of *M. p. aurita* has remained comparatively poorly understood.

Recently, Wangmo, et al. ^27^ describe a new species, *Manis indoburmanica*, from South Tibet, which highlights several challenges in modern taxonomy. The recognition of this new species is based solely on the mitochondrial genome, and lacks a morphological comparison with *M. pentadactyla*, thus failing to establish a clear morphological boundary for *M. indoburmanica*. Critically, as noted by Zijlstra ^32^, Wangmo, et al. ^27^ did not evaluate whether their samples correspond to existing historical names like *M. p. aurita* ^33^ or *M. assamensis* ^34^, leaving the novelty of the taxon in question. Such challenges underscore the necessity of integrative taxonomic approaches that incorporate genome-wide data, robust morphological comparisons with both *M. pentadactyla* and *M. p. aurita* ^35–37^.

To resolve this taxonomic uncertainty, we employed an integrative taxonomic framework combining genomic and morphological evidence ^38^, incorporating the type specimens of *M. p. aurita*. Species delimitation was evaluated under a phylogenetic diagnostic species concept (dPSC), which operationally recognizes species as reciprocally monophyletic and diagnosable lineages ^39^. The objective of this study was to evaluate whether pangolins from South Himalaya correspond to the subspecies *aurita* and, if so, whether their degree of genetic and phenotypic distinctiveness warrants recognition at full species rank. Our analyses reveal clear and pronounced differentiation from *M. pentadactyla* across both genetic and multiple morphological datasets, providing robust support for recognizing this lineage as the distinct species *Manis aurita* Hodgson, 1836.

## RESULT

### Genetic evidence for the distinctiveness of *M. aurita*

We generated seven new NGS datasets for *Manis aurita*, including the *M. aurita* lectotype (BMNH 43.1.12.85) (**Tables S1, S2**), and integrated them with previously published data. The final comprehensive dataset for genomic analysis comprised 55 whole-genome sequencing (WGS) datasets (representing 53 *Manis* individuals, one *Smutsia gigantea*, and one *Phataginus tricuspis;* (**Table S3**) and 70 mitogenomes (**Table S2**). Phylogenetic reconstruction based on autosomal SNPs and mitogenomes yielded consistent and robustly resolved Maximum Likelihood (ML) trees, with maximal bootstrap support (bootstrap support [BS] = 100) for all interspecific relationships (**Figure 1A**). The topology corroborates all previous findings ^10,13,14,18,27^, identifying *Manis pentadactyla* sensu lato (including all three subspecies *pentadactyla*, *pusilla*, and *aurita*) as the sister clade to all other Asian congeners. The putative taxon “mysteria” emerges as the immediate sister to *M. javanica.* Within *M. pentadactyla* s. l. (n=44), samples clustered into two deeply diverged clades, corresponding to MPA and MPB identified in Hu, et al. ^13^. Finally, sex-chromosome-derived SNPs, and single-copy orthologues (**Supplementary Figures S1**, **S2**) also yielded topologies identical to the autosomal SNP and mitochondrial trees, reinforcing these phylogenetic relationships.

**Figure 1:**
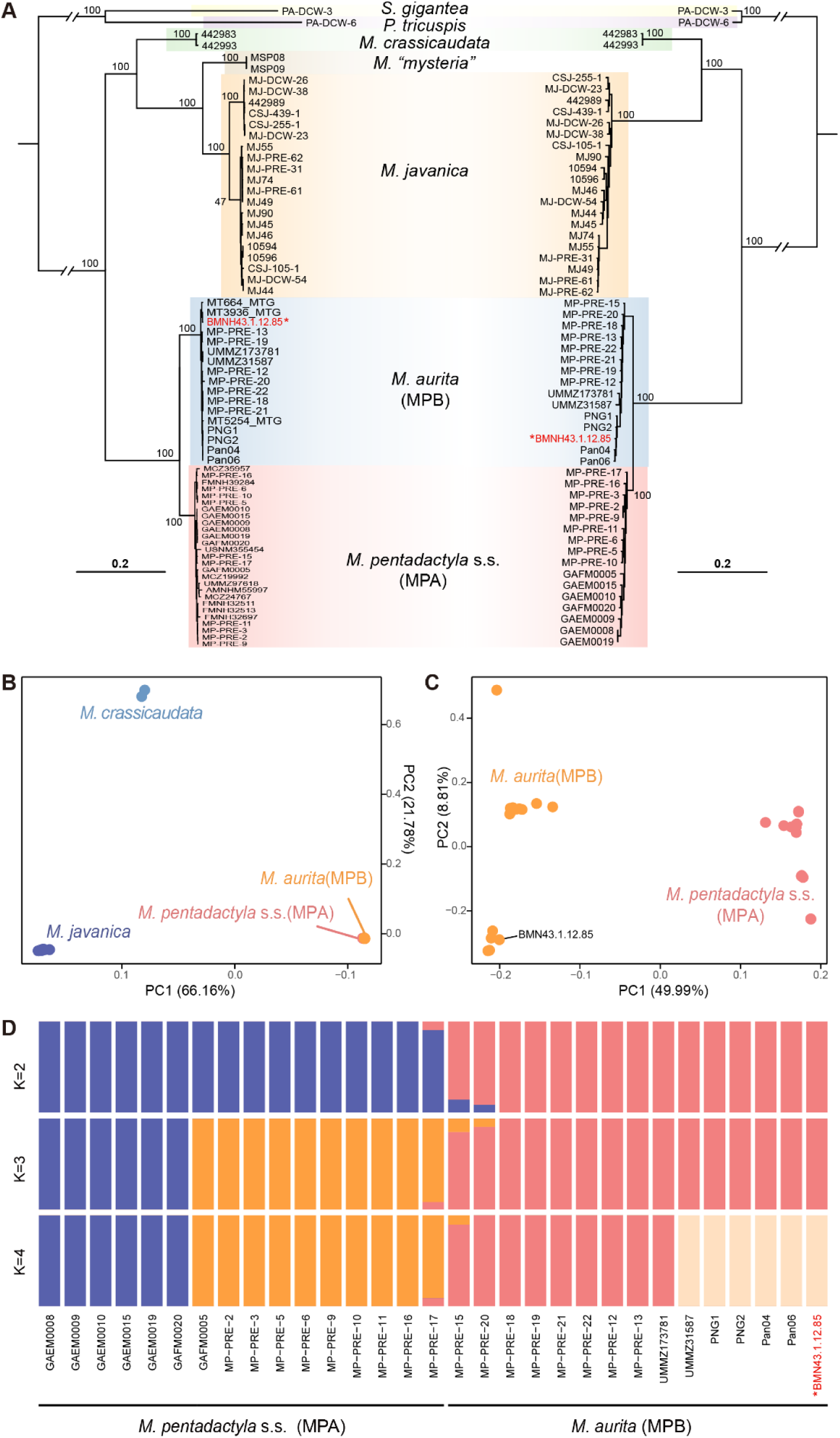
Phylogenomic relationships and population structure of pangolins. (A) Maximum likelihood (ML) phylogenies based on complete mitochondrial genomes (left) and autosomal nuclear single-nucleotide polymorphisms (SNPs) (right). Numbers at nodes indicate bootstrap support values. Scale bars represent substitutions per site. (B) Principal Component Analysis (PCA) of genomic SNP data demonstrating the separation of four *Manis* species. (C) PCA restricted to the *M. pentadactyla* s.l. illustrating clear genetic differentiation between *M. pentadactyla* s.s. (MPA) and *M. aurita* (MPB). Values in parentheses indicate the percentage of total variance explained by each principal component. (D) Population structure of the *M. pentadactyla* species complex inferred by ADMIXTURE assuming K=2 to K=4. Each vertical bar represents an individual, with colored segments denoting the proportion of ancestry from each inferred cluster. The lectotype of *M. aurita* is marked with a red star.

When SNP of *Manis* species were projected onto principal-component (PC) space, every *M. pentadactyla* s. l. sample falls into a tight cluster that is clearly distinct from the other congeneric species (**Figures 1B**, **1C**). Restricting the analysis to *M. pentadactyla* s. l. reveals non-overlapping splitting between MPA and MPB along PC1 (Tracy-Widom test, p = 7.49 × 10^-6^; **Table S4**), underscoring consistent genomic differentiation despite their close relationship. The result of ADMIXTURE analysis echoes these patterns (**Figure 1D**). The lowest cross-validation error occurs at K = 2 (CV = 0.47), with marginal increases at K = 3 (CV = 0.50) and K = 4 (CV = 0.53; **Table S5**). At K = 2, the inferred clusters correspond closely to MPA and MPB, with three individuals displaying minor ancestry from the opposite cluster. Of particular note, in all above phylogenetic and population genetic analyses, the lectotype of *M. aurita* fell in the MPB clade. Taken together, the concordant mitochondrial and nuclear results robustly validate that MPB represents *M. aurita* and is genetically distinct from the *M. pentadactyla* sensu stricto (MPA) (which represent the other two subspecies).

To explore the geographic distribution of *M. aurita*, we augmented our mitogenome dataset with publicly available georeferenced mitochondrial *CYTB* and *COI* sequences representing *M. pentadactyla* s.l.. This expanded phylogenetic analysis recovered an identical backbone topology (Figure 2A). Critically, samples attributed to *Manis indoburmanica* (from South Tibet) were robustly embedded within MPB (BS=100). Furthermore, mapping the occurrences of MPB haplotypes revealed a distribution confined to Nepal, South Tibet, and Assam (Figure 2B). Collectively, these lines of evidence demonstrate that *M. indoburmanica* is conspecific with *M. aurita* and therefore represents a junior synonym of this species, whose distribution is restricted to the southern Himalayas.

**Figure 2:**
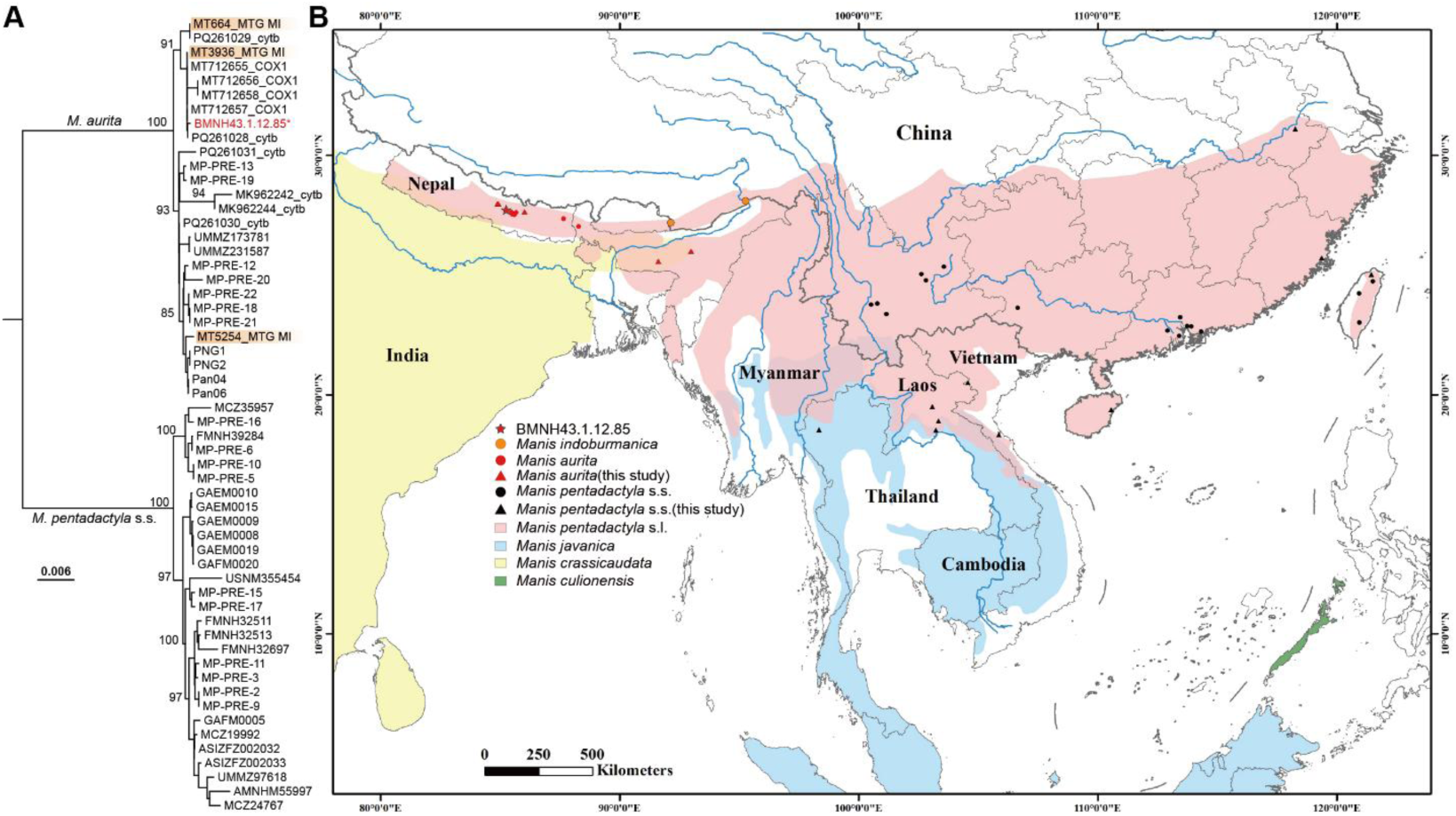
Mitochondrial phylogeny and geographic distribution of the *Manis pentadactyla* species complex. (A) Maximum likelihood phylogeny constructed from an expanded dataset of mitochondrial sequences (complete mitogenomes and partial *CYTB* and *COX1* fragments), integrating the new sequences generated in this study with publicly available data. Numbers at the nodes represent bootstrap support values. The scale bar indicates the number of substitutions per site. *M. indoburmanica* specimens (MI, highlighted in orange color) and lectotype of *M. pentadactyla aurita* (BMNH 43.1.12.85) nest firmly within the *M. aurita* (MPB) clade. (B) Geographic distribution map with verified locality information (**see Table S1**). Circles represent data obtained from previous studies while triangles indicate samples collected in this study. Shaded areas denote the approximate ranges of Asian pangolin species according to IUCN Red List.

### Morphometric evidence for the distinctiveness of *M. aurita*

To assess whether *M. aurita* is morphologically distinct from the *M. pentadactyla* sensu stricto (including subspecies *pentadactyla* and *pusilla*) and thus merits recognition as a separate species, we employed linear measurements-based morphometric and three-dimensional landmark-based geometric morphometric analyses. PCA of 20 cranial measurements (**Figure S3A**) for 44 specimens positioned *M. pentadactyla* s. l. within a morphospace that showed partial overlap with the Indian pangolin (*M. crassicaudata*) but remained clearly separated from the Sunda (*M. javanica*) and Philippine (*M. culionensis*) pangolins (Figure 3A; **S4A**, **S4B**). Along PC1, which accounted for 39.4% of the shape variance and primarily reflected overall cranial size, specimens identified as *M. aurita* consistently occupied the positive extreme. This placement indicates that *M. aurita* possesses a significantly larger skull, and proportionally shorter palatine when compared to *M. pentadactyla* s.s. (**Tables S7A-6C**).

**Figure 3:**
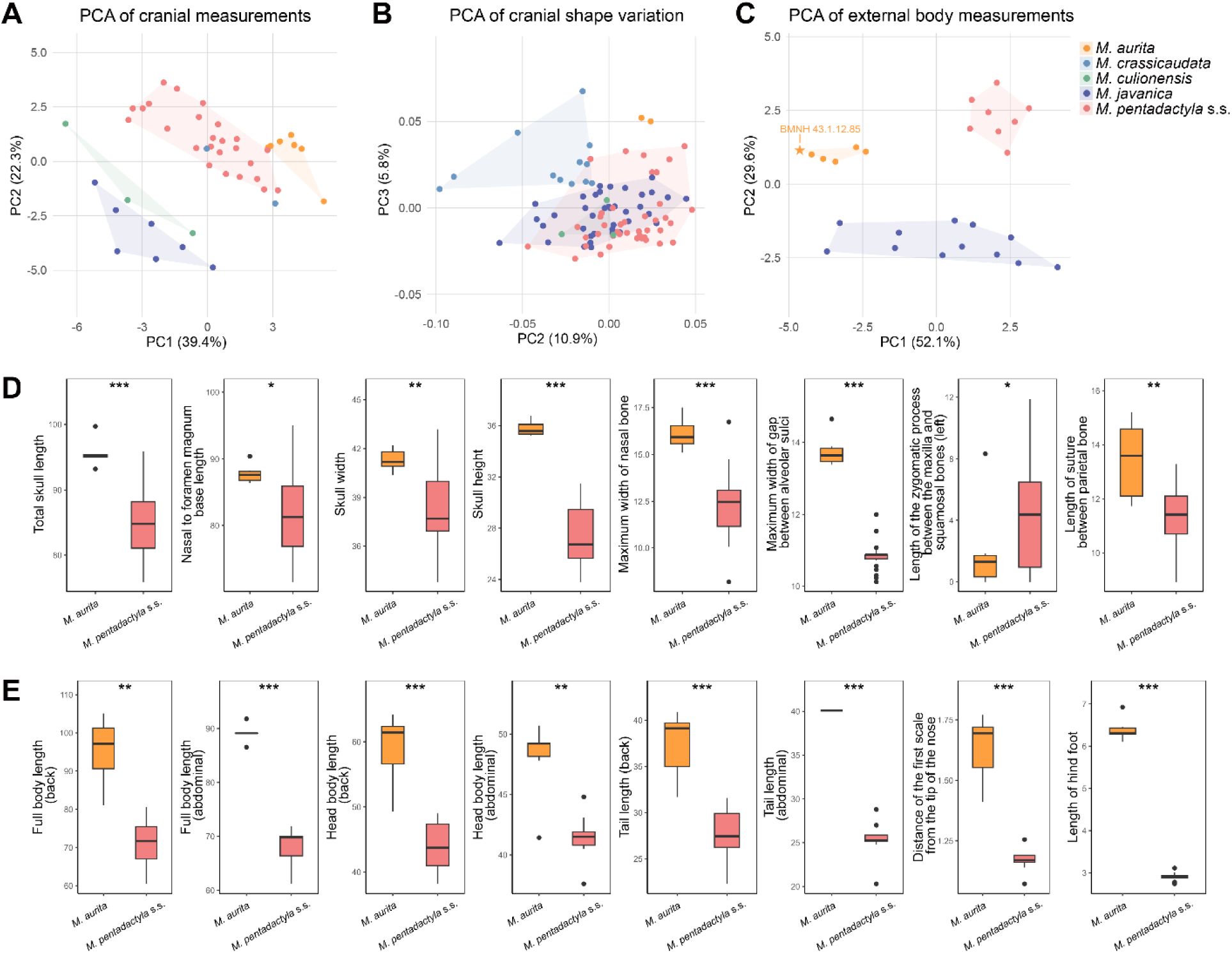
Morphometric analyses based on cranial and external morphology. (A) Principal Component Analysis (PCA) based on 20 linear cranial measurements (PC1 *vs*. PC2). (B) PCA of cranial shape variation derived from 75 3D geometric morphometric landmarks (PC2 *vs*. PC3). (C) PCA based on 14 external body measurements (PC1 *vs*. PC2). For all PCA plots (A-C), values in parentheses on the axes indicate the percentage of variance explained. Note the inclusion of the *M. aurita* lectotype (BMNH: 43.1.12.85) in panel C, which clusters firmly with other *M. aurita* specimens. (D, E) Boxplots comparing key cranial (D) and external (E) measurements between the two taxa. Boxes depict the interquartile range (IQR) with the median indicated by the central line; whiskers extend to 1.5 times the IQR. Individual dots indicate outliers beyond 1.5 × IQR. Units are in millimeters (D) and centimeters (E). Statistical significance from Student’s t-tests is denoted as: * *P* < 0.05; ** *P* < 0.01; *** *P* < 0.001. See Tables S5–S14 and Figures S4, S10, S11 for additional results.

Three-dimensional landmark-based geometric morphometric analyses of 75 cranial landmarks (**Table S6; Figure S5**) obtained from micro-CT scans successfully distinguished *M. aurita* from *M. pentadactyla* s.s. along PC3 (Figure 3B; **Tables S8A**, **S8B**), but not PC1 or PC2 (**Supplementary Figures S4C**, **S4D**). Compared to *M. pentadactyla* s.s., *M. aurita* exhibits a relatively shorter yet broader nasal bone, a laterally expanded junction between the maxilla and palatine bones, a posteriorly positioned anterior point of the foramen magnum on the ventral occipital surface (landmark 25, **Figure S5**; **Table S6**), and an anteriorly shifted M-shaped suture between the maxilla and palatine bones on the ventral view, culminating in a shorter ventral maxilla (**Supplementary Figure S10C**).

We also conducted a PCA based on fourteen external body measurements (**Figure S3A**) for *M. aurita, M. pentadactyla* s.s. and *M. javanica.* A crucial aspect of this analysis was the inclusion of the *M. aurita* lectotype, which notably lacks skull material. The results demonstrated that *M. aurita* separated from *M. pentadactyla* s.s. along both PC1 (accounting for 52.1% of variance) and PC3 (6.9% of variance) (Figure 3C; **Supplementary Tables S9A-9C**, **Figures S4E**, **S4F**). The PCA-based separation is strongly supported by *t*-tests on individual cranial (Figure 3D; **Supplementary Figure S6A**; **Table S10**) and external measurements (Figure 3E; **Supplementary Figure S6B**; **Supplementary Table S11**), which show *M. aurita* having significantly longer body lengths (89.14±1.65 cm; n=6), back tail lengths (37.39±3.73 cm; n=6), larger hind feet (6.39±0.28 cm; n=6), an extended snout and markedly smaller ears (2.01± 0.05 cm; n=6) (**Supplementary Data S1**).

Finally, Canonical Variate Analyses (CVAs), using both cranial and external variables achieved 100% classification accuracy in separating *M. aurita* and *M. pentadactyla* s.s. (**Tables S12**, **S13, S14**; **Supplementary Figure S7**; **Supplementary Data S2**). In contrast to these clear differences in skeletal and external body proportions, an analysis of scale characteristics using superimposed, scale-encoded heatmaps of 12 intact body skins (nine *M. pentadactyla* s. s. and *three M. aurita*) revealed no significant differences in either the total number of scale rows or their spatial distribution between the two taxa (**Supplementary Figure S8**; **Supplementary Table S15**; **Supplementary Data S3**, **S4**). Collectively, the consistent and pronounced size- and shape-based differences in both cranial and external morphology, strongly demonstrate that *M. aurita* is morphologically distinct from *M. pentadactyla* s.s. supporting its full species status.

### Evolutionary and demographic history of *Manis aurita*

We estimated the divergence time within pangolins using 200 selected clock-like single-copy orthologous genes. The most recent common ancestor (MRCA) of the genus *Manis* was dated to approximately 29.4 million years ago (Ma) (95% confidence interval [CI] = 27.4 to 31.5 Ma; Figure 4A), a finding consistent with Gu et al. (2023). *M. aurita* (MA hereafter) and *M. pentadactyla* s. s. (MP hereafter) diverged approximately 1.8 Ma (95% CI = 1.3 to 2.5 Ma), during the early Pleistocene.

**Figure 4:**
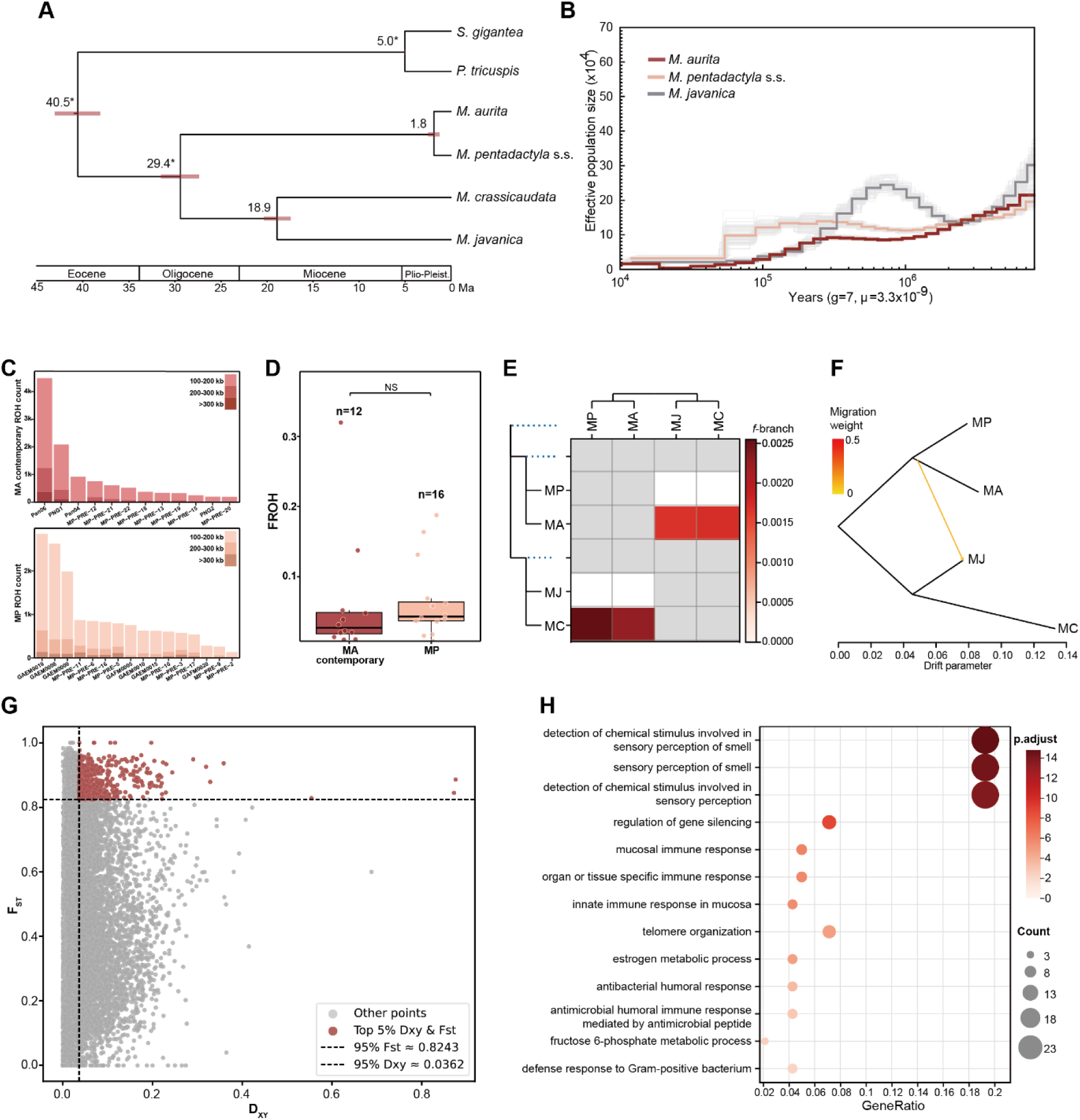
Divergence time, demographic dynamics, and genomic divergence of *Manis* species (A) Divergence time estimates for pangolins based on 200 clock-like single-copy orthologous genes, inferred using *BEAST3. Numbers at nodes represent mean age estimates in millions of years (Ma), while the associated horizontal bars represent the 95% Confidence Intervals (CI). Asterisks indicate nodes used for calibration. (B) Historical effective population size (Nₑ) trajectories for three species inferred using PSMC. Shaded areas represent 100 bootstrap replicates. (C-D) Comparison of recent inbreeding levels between the contemporary population of *M. aurita* (MA contemporary) and *M. pentadactyla* s.s. (MP): (C) Distribution of Runs of Homozygosity (ROH) across individuals in MA contemporary and MP and (D) Genome-wide fraction of ROH (F_ROH_). Differences between groups were assessed using a Wilcoxon rank-sum test; no statistically significant difference was detected (P > 0.05). (E-F) Interspecific gene flow analyses among four *Manis* species. MC: *M. crassicaudata*, MJ: *Manis javanica*: (E) Heatmap of Dsuite f-branch statistics, where warmer colors indicate stronger signals of admixture on a given branch. (F) TreeMix population graph showing inferred drift and a migration event (arrow). (G-H) Genomic differentiation between *M. aurita* and *M. pentadactyla* s.s. (G) Manhattan plot of F_ST_ and D_XY_ values, with dashed lines indicating the 95% outlier thresholds used to define genomic islands of divergence. (H) Gene Ontology (GO) enrichment analysis of genes within the identified genomic islands, revealing significant enrichment for terms related to olfactory functions. Dot size corresponds to gene count and color to statistical significance (p.adjust).

Pairwise Sequentially Markovian Coalescent (PSMC) analyses revealed different demographic trajectories for the two species (Figure 4B). Following an initial post-divergence decline, the MP population remained relatively stable from 1.0 Ma until a pronounced reduction approximately 50 ka. In contrast, the MA population exhibited demographic stability until ∼300 ka, after a sustained long-term decline.

We employed GONE to estimate N_e_ fluctuations over the most recent 600 generations. For MP, a sharp population contraction was detected approximately 60–50 generations ago (corresponding to the 17^th^ century), during which N_e_ plummeted to 2606 (ranging from 2387 to 2966). By comparison, MA experienced an earlier bottleneck approximately 100-110 generations ago (corresponding to the 14th century), with N_e_ declining to 1802 (1648-2033) (**Supplementary Figure S9**).

Given the inferred population declines and reduced contemporary N_e_ in both species, we evaluated whether these demographic changes were accompanied by increased inbreeding. ROH profiles varied substantially among individuals (Figure 4C). In both contemporary MA and MP, short ROHs (100–200 kb) constitute the majority. However, a subset of individuals showed conspicuously elevated counts of longer ROHs (200–300 kb and especially >300 kb), consistent with more recent parental relatedness. Although the mean F_ROH_ is slightly higher in MP (n = 16; ∼0.063) than in contemporary MA (n = 12; ∼0.060), the difference is not significant (Wilcoxon rank-sum test, P > 0.05). Notably, several outliers in both species exhibit exceptionally high F_ROH_ (> 0.1), including MA individuals Pan06 and PNG1 and MP individuals GAEM0008, GAEM0009, and GAEM00019, indicating pronounced, localized inbreeding in particular populations, such as MA around the Kathmandu Valley (Figure 4D)

We subsequently examined patterns of interspecific gene flow involving MA to assess the impact of current and historical range overlap with its congeners (Figure 2B). D-suite analyses revealed significant, albeit low-level, introgression from the Indian pangolin (*M. crassicaudata*) into both MP (f4-ratio ≈ 0.0025, p < 1×10-15) and MA (f4-ratio ≈ 0.0022, p < 1×10-15) (Figure 4E; **Supplementary Table S16**). We also detected subtle genomic contributions from MA to the Malayan pangolin (*M. javanica*) supported by D-suite (f4-ratio ≈ 0.0017) and TreeMix (Figure 4F). Fastsimcoal2 strongly supported absolute reproductive isolation between MP and MA post-divergence (ΔAIC > 2.9; **Supplementary Table S17**). Collectively, these results indicate that MA has experienced minimal introgression from other pangolin species, and has maintained profound and near-complete genetic isolation from its sister species, MP, since their divergence.

Finally, we compared MA and MP genomes for signatures of local adaptation by calculating nucleotide diversity (π) and fixation index (F_ST_), and absolute genetic differentiation (D_XY_) (**Supplementary Figure S10**). Genomic islands of divergence were defined as windows in the top 5 % for both F_ST_ (>0.8575) and D_XY_ (>0.0373; Figure 4G). Gene Ontology (GO) enrichment analysis of genes within these identified islands revealed significant enrichment for olfactory-related functions, including “the detection of chemical stimuli and broader sensory perception of smell” (Figure 4H).

## DISCUSSION

### The Revalidation of *M. aurita*

Despite accumulating molecular evidence suggesting overlooked, or cryptic species ^13,14,18^, the taxonomy of *M. pentadactyla* s. l. lacked rigorous examination. Our mitochondrial and genome-wide phylogenomic analyses consistently reveal a divergence between *M. aurita* and *M. pentadactyla* s.s. at about 1.8 Ma (Figure 4A). The magnitude of this split surpasses commonly accepted thresholds for intraspecific variation ^40–42^. Complementing this genetic divergence, morphometric analyses identify consistent and diagnostic differences that distinguish *M. aurita* from *M. pentadactyla* s.s. (Figure 3). Cranially, *M. aurita* possesses a significantly larger overall skull with a proportionally shorter and broader nasal region compared to *M. pentadactyla* s.s. (**Supplementary Figures S11**; **Supplementary Tables S9**, **S10**). Externally, *M. aurita* is distinguished by larger body size, a longer tail and smaller pinnae compared to *M. pentadactyla* s.s. (**Supplementary Figures S12)**.

The distribution of *M. aurita* is associated with the Himalayan region (Figure 2B). It includes the type locality of *M. aurita* from central Nepal, as well as individuals from South Tibet recently described as *Manis indoburmanica* ^27^. In contrast, *M. pentadactyla* s.s. is predominantly distributed in South and Eastern China and Southeast Asia. Our results suggest that their deep divergence (∼1.8 Ma) was initiated during Pleistocene climatic oscillations and was followed by pronounced population declines, with each lineage likely persisting in fragmented habitats within isolated refugia.

Although pangolins have been reported to occur locally up to ∼3,000 m asl and are capable of swimming ^43^, they typically occupy forested habitats at lower elevations (<2,500 m), and documented swimming behavior largely involves still or slow-moving water. Therefore, while no single mountain or river acts as an absolute barrier, the synergy of geographic and ecological filters has effectively terminated gene flow. Major landscape features, such as the Yarlung Tsangpo–Brahmaputra drainage ^6,44^ and the rugged Arakan Mountains ^45^, combined with steep climatic and vegetational gradients, have enforced long-term isolation. This pattern of “synergistic filtering” mirrors the evolutionary history of various other Himalayan mammals and herpetofauna ^6,44–50^. This sustained isolation has ultimately driven speciation. However, to precisely define the phylogeographic boundaries and identify potential contact zones between *M. aurita* and *M. pentadactyla*, focused sampling across the data-deficient regions of northern Myanmar remains essential.

Collectively, the congruent evidence from our genetic, morphological, and distributional analyses clearly supports recognizing Hodgson’s *aurita* as a distinct species. While Zijlstra ^32^ has argued for its revalidation, our study substantiates this view with broader and more integrative evidence. Consequently, in accordance with the Principle of Priority of the International Code of Zoological Nomenclature (ICZN), we reinstate *Manis aurita* Hodgson, 1836 as the valid name for this species. This action renders junior synonyms such as *Pholidotus assamensis* Fitzinger, 1872, and *Manis indoburmanica* Wangmo, Ghosh, Dolker, Joshi, Sharma and Thakur, 2025.

### Evolutionary history of *M. aurita*

The population genetic and demographic analyses illuminate the evolutionary trajectory of *M. aurita* following its speciation. The ancestors of *M. aurita* and *M. pentadactyla* diverged in the early Pleistocene, period of, significant global climatic oscillations. Given that pangolins are generally warm-adapted species, with no historical fossil evidence suggesting their presence in northern China ^51^, the climatic shifts likely played a crucial role in driving the allopatric separation of the ancestor into western (Himalaya) *vs*. eastern (East/ Southeast Asia) refugium ^46–50^. Pangolins are highly olfactory-reliant mammals, exhibiting significantly enlarged olfactory bulb and turbinals ^52–54^, using smell to forage for ants and termites ^55^. As *M. aurita* and *M. pentadactyla* adapted to their respective environments, certain OR genes likely diverged under selection as supported in our GO enrichment analysis (Figure 4H).

Our results highlight contrasting demographic trajectories between M. aurita (MA) and M. pentadactyla s.s. (MP) driven by regional climatic variations. The stability of MP’s N_e_ until ∼50 ka underscores the role of eastern and southern China’s lowland forests as stable refugia that provided substantial climatic buffering ^56^. Conversely, MA underwent a sharp demographic collapse starting ∼300 ka, triggered by the intense cooling and aridification of the Penultimate Glaciation ^57,58^. This pattern mirrors the stress-induced declines seen in other high-altitude taxa, such as snub-nosed monkeys ^59^. While the maritime-influenced climate of southern China mitigated glacial impacts for MP ^60^, the restricted range and severe habitat instability of the Himalayan region rendered MA uniquely susceptible to long-term decline.

Interestingly, the recent demographic declines inferred by GONE analysis for both species align temporally with the Little Ice Ages (LIAs). The 17^th^-century bottleneck observed in MP coincides with the well-known “Late Ming Little Ice Age” in China. This era was characterized by severe cooling and megadrought ^61,62^ which has been linked to agricultural failure and the eventual collapse of the Ming Dynasty ^63^. Similarly, the demographic contraction of MA around the 14th century corresponds to the onset of the LIA in the Himalayan region. While direct documented evidence linking these hydroclimatic perturbations to contemporaneous sociopolitical transitions in the Himalaya remains limited and debated, multiple independent proxy archives indicate pronounced hydroclimate volatility during the early LIA, including multi-decadal droughts as well as intervals of enhanced precipitation ^64,65^. Our findings underscore that even relatively recent, short-term climatic oscillations like the LIA can leave a discernible and lasting footprint on the genomic diversity of pangolins.

Our analyses further detected limited gene flow between *M. aurita* and two other Asian pangolins. The signal of introgression between *M. aurita* and *M. crassicaudata* (contributing to <0.25% of the genome) is unsurprising, given their sympatric ranges in southern Nepal and northeastern India. We also observed a similarly low level of gene flow between *M. aurita* and *M. javanica*, indicating that their distributions likely have overlapped at least historically. Despite these instances of historical admixture, the low level of gene flow indicates *M. aurita* has maintained its status as a distinct evolutionary entity.

It is noteworthy that in the admixture analysis, at both K=3 and K=4, *M. aurita* separated into two distinct genetic clusters: one comprising individuals from Nepal and the other comprising individuals from South Tibet and northeastern India, a pattern that indicates significant intraspecific subdivision (Figure 1D).

### Conservation Implications

The formal recognition of *M. aurita* as a distinct species carries profound conservation significance. Its intrinsic vulnerability is underscored by a geographically restricted range, confined to the Himalayan–Indo-Burmese region, and a demographic history marked by a more pronounced and sustained population decline since the Middle Pleistocene compared to *M. pentadactyla*.

Despite the long-term population decline revealed by PSMC analyses, *M. aurita* on average exhibits lower levels of recent inbreeding than *M. pentadactyla* as evidenced by the results of ROH, F_ROH_ (Figures 4C, **4D**) and GONE (**Supplementary Figure S9**). This likely reflects differing histories of anthropogenic pressure nowadays; the less intensively developed landscapes of South Asia historically provided more contiguous refuge habitats compared to the range of *M. pentadactyla* in China and Southeast Asia, where has experienced intense human activity for centuries, leading to severe habitat fragmentation and the regional extinction of many medium-to-large mammal species ^66–68^. However, it is noteworthy, *M. aurita* around Kathmandu valley exhibit extremely high inbreeding levels (i.e., F_ROH_ > 0.1), indicating that these populations are already facing severe inbreeding depression (Figure 4D). Conservation strategies must therefore prioritize the identification and management, and continue search for other locally vulnerable populations to prevent local extinctions.

Overarching these intrinsic vulnerabilities is the critical threat from illegal wildlife trade. Genetic analyses of confiscated scales have repeatedly confirmed that *M. aurita* is a target ^13^. Alarmingly, our findings also reveal that products from this species have infiltrated regulated traditional medicine markets (data from Xie, et al. ^69^; **Supplementary Figure S13**), demonstrating that illegally sourced materials are being laundered through formal supply chains. The comprehensive genomic dataset presented here, with its improved geographic coverage and inclusion of previously unsampled populations, provides an enhanced genetic baseline which will bolster wildlife forensics by allowing for greater precision in determining the geographic provenance of confiscated scales, thereby helping to pinpoint poaching hotspots.

The recognition of *M. aurita* as a distinct species has immediate implications for international regulation and enforcement. All eight currently recognized pangolin species were transferred to CITES Appendix I, thereby prohibiting international commercial trade in those named taxa and their parts under the Convention. We encourage timely coordination between taxonomic updates and regulatory instruments. Specifically, rapid incorporation of revalidated taxa into the standard nomenclatural references used by CITES, and inclusion of *M. aurita* under Appendix I.

### Taxonomy

*Manis aurita* Hodgson, 1836 stat. nov.

*Manis auritus* Hodgson, 1836. Original description.

*Manis aurita* Waterhouse,1838 *Pholidotus assamensis*

Fitzinger, 1872 *Manis pentadactyla auritus* Thomas 1918

*Manis pentadactyla* Ellerman & Morrison-Scott, 1951 (part)

*Manis indoburmanica* Wangmo, Ghosh, Dolker, Joshi, Sharma and Thakur, 2025

Lectotype: BMNH 43.1.12.85, an adult (skin) collected by Brian Houghton Hodgson in 1836 from Central Nepal (**Supplementary Figure S12F**).

### Emended diagnosis

*Manis aurita* is a medium-sized Asian pangolin from the South Himalaya; with a total length longer than *M. pentadactyla* but shorter than *M. crassicaudata.* The species is characterized by a flap-shaped external pinna that is smaller than *M. pentadactyla* (**Supplementary Figures S11A**, **S11E-F)**. The rostral scales do not reach the tip of the snout, leaving a distinct patch of exposed, naked skin (typically connected to the nasal tip in *M*. *crassicaudata and M. culionensis*) (**Supplementary Figure S11A**). The dorsal and ventral scales of *M. aurita* are uniformly dark brown to deep chestnut (overall yellowish-brown in *M. crassicaudata* and pale yellow on the abdomen and distal limbs in *M. javanica*). The medial rows of dorsal scales are broadly trapezoidal with straight posterior margins (typically broad rhombic shape in *M. pentadactyla* and *M*. *crassicaudata*, and elongated rhombic and sharply pointed in *M*. *javanica* and *M. culionensis*) (**Supplementary Figure S11B, Table S18**).

Cranially, *M. aurita* is distinguished by a shorter palatal process of the premaxilla (extending less than half the premaxillary length), in contrast to *M. crassicaudata*, and *M. culionensis*, where the process exceeds half the length (**Supplementary Figures S11C, S12B**). The incisive foramen is oval-shaped and enclosed entirely within the premaxilla, neither extending to the posterior border nor contacting the ventral projection of the maxilla, whereas in *M. crassicaudata* and some specimens of *M. javanica*, the incisive foramen is formed by the premaxilla (anteriorly/laterally) and the maxilla’s palatine processes (posteriorly). The maxillae in *M. aurita* are inflated, especially anteriorly where it meets the premaxilla, resulting in a more robust rostral profile; by contrast, in the other four *Manis* species, the rostral region appear slenderer (**Supplementary Figures S11C**, **S12A, Table S18**).

In M. aurita, the zygomatic arch is complete or nearly so, representing a plesiomorphic state ^70^ (**Supplementary Figure.S11D**). This contrasts sharply with *M. crassicaudata*, where the arch is incomplete, and *M. culionensis*, in which it is consistently absent. The postglenoid foramen opens on the lateral aspect of the squamosal zygomatic process (apomorphic; see <u>Gaudin and</u> Wible ^70^ whereas it opens on the posterior aspect in *M. javanica* and *M. culionensis*) **(Supplementary Figures S11D, S12D).** The jugular and hypoglossal foramina open within a single depressed fossa (apomorphic; whereas in the other four *Manis* species, these foramina are not recessed) (**Supplementary Figures S11C**, **S12C**).

### Description

*Manis aurita* has an elongated head, short limbs, and a tail shorter than half the body length. Its external pinna is small and flap shaped. The head and body are covered in overlapping, exceptionally smooth keratin scales: dorsal scales are broadly trapezoidal with straight posterior margins. The lateral abdominal scales are broad and shield shaped consistently observed across all Asian pangolins^30,70^. On the snout, the first row of head scales lies well behind the nasal tip, leaving a bare patch of skin. Scale coloration in adult’s deep chestnut dark brown; juveniles are slate-grey or greyish brown, with more bristles between scales.

The skull presents a robust overall profile. The rostrum is notably thickened due to the inflated maxillae, particularly at their junction with the premaxilla. The ventral palatal process of the premaxilla is short, extending less than half the length of the premaxilla. The incisive foramen is oval and situated entirely within the premaxilla, without reaching the posterior border or contacting the ventral projection of the maxilla. In ventral view, the vomer is not visible. From a dorsal perspective, the nasal bones are broad and extend posteriorly to the parietal, terminating in an arch-like shape and sharing a broad suture with the maxilla. The braincase is dorsally inflated, resulting in a dome-shaped appearance.

The postglenoid foramen opens laterally on the zygomatic process of the squamosal, a condition recognized as an apomorphy for Pholidota ^70^. In the zygomatic region, the maxillary process either connects to the squamosal via an ossified, rod-like structure or remains detached—an apomorphic trait in certain lineages ^70^. The pterygoid does not extend beyond the posterior margin of the tympanic bulla. The jugular and hypoglossal foramina open within a single depressed fossa on the ventral surface of the cranium. External and cranial measurements are provided in Supplementary Data S1 and S2, respectively.

The total scale count ranges from 538 to 596. The trunk, spanning from the neck to the tail base, bears 254–274 scales arranged in 23–24 longitudinal and 15–17 transverse rows. The tail (caudal) scales total 140–153, 80–94 on the dorsal surface organized into five transverse by 16–20 longitudinal rows, and 50–63 on the ventral surface in three to five transverse by 18–20 longitudinal rows, with 17–19 scales along each lateral edge. Unilateral scale counts for the forelimbs and hindlimbs range from 37–55 and 32–41, respectively.

### Comparisons

MA is distinguished from its closest relative, MP, by its larger size (average total length = 95.2 cm *vs*. 71.2 cm), relatively longer tail (average 37.4 cm *vs*. 27.5 cm), and significantly larger hindfoot (average LHF = 6.4 cm *vs*. 2.9 cm), and distinctly smaller pinnae (average 2.01 cm *vs* 2.18 cm) (Figure 3E; **Supplementary Figures S6B**, **S11E–F**).

The skull of MA is larger and taller than MP (average cranial width = 95.6 mm *vs*. 84.6 mm; average height = 35.8 mm *vs*. 27.6 mm). It can be distinguished from MP by the combination of the following characters: shorter and broader nasal bones (length-to-width ratio c. 1.9:1 *vs*. 2.5:1 in MP); a more robust rostral profile due to anteriorly inflated maxillae that bulge laterally (*vs*. slender in MP); and a larger distance between alveolar sulci (average = 13.8 mm *vs*. 10.9 mm) (Figure 3D; **Supplementary Figure S6A**). The palatine bone is positioned more anteriorly in MA, with the most posterior point of the palatine suture aligning with the junction of the maxillary and squamosal zygomatic processes, whereas it lies posterior to this junction in MP (**Supplementary Figures S12A**). Uniquely among Asian pangolins, the jugular and hypoglossal foramina in MA open within a single depressed fossa on the ventral surface of the cranium, while they open separately and are not recessed in all other species (**Supplementary Figure S12C**).

*Manis aurita* is easily distinguished from *M. javanica* by its dark brown/chestnut coloration (*vs*. pale yellow restricted to the distal limbs, abdomen, and the posterior half of the tail in *M. javanica*), its trapezoidal dorsal scales (*vs*. sharp/pointed), and by the postglenoid foramen opening on the lateral aspect of the squamosal zygomatic process (*vs*. posterior aspect in *M. javanica*) (**Supplementary Figure S12D**).

MA differs from *M. culionensis* by possessing trapezoidal dorsal scales (*vs*. sharp/pointed) and a complete zygomatic arch (*vs*. consistently absent). Cranially, it is distinguished by a laterally opening postglenoid foramen (*vs*. opening on the posterior aspect), a shorter palatal process of the premaxilla (extending less than half the premaxillary length *vs*. exceeding half), and closer proximity of the maxillary and squamosal zygomatic processes (**Supplementary Figure S12B, S12D**).

Finally, it is distinguished from *M. crassicaudata* by its smaller body size, larger external ear, complete zygomatic arch (*vs*. consistently absent ^30^), incisive foramen enclosed entirely within the premaxilla (*vs*. formed by premaxilla and maxilla), and absence of a visible vomer in ventral view (*vs*. visible vomer in part of the specimens) (**Supplementary Figure S12B**).

### Distribution, habitat, and Conservation Statues

*Manis aurita* is distributed along the southern Himalayan foothills, with confirmed occurrences in Nepal, South Tibet and northeastern India (including Assam), southern Bhutan, and extending into southern Tibet. The species’ altitudinal range spans from lowland areas to approximately 3,000 meters above sea level ^71,72^. The eastern extent of its distribution remains undefined, but the Arakan Mountains appear to represent the eastern geographic limit for this species ^45^. Further surveys are necessary to ascertain its potential presence in Myanmar.

*Manis aurita* predominantly inhabits mixed deciduous and evergreen forests. Key tree species in these habitats include *Shorea robusta, Schima wallichii, Castanopsis indica*, and *Alnus nepalensis* ^73^ ^74^. *M. aurita* is also frequently observed in forested areas adjacent to human-modified landscapes, indicating some level of adaptability to disturbed environments ^72–75^.

Despite protected under national laws such as the Nepal’s National Parks and Wildlife Conservation Act (1973), India’s Wildlife Protection Act (Schedule I, 1972), and internationally under CITES Appendix I, *M. aurita* remains a significant target for illegal wildlife traffickers in South Asia. For example, Kathmandu functions as a major transit hub for smuggled pangolin scales ^19,21^. Similar trafficking pressures are evident in neighboring India and Bhutan ^20^, where challenges such as limited enforcement capacity and porous international borders facilitate cross-border smuggling operations.

## MATERIAL AND METHODS

### Taxon sampling and sequencing

DNA was extracted from tissue samples (n=21) and purified using the Dneasy blood and tissue kit (Qiagen). Genomic DNA was sheared to ∼350 bp fragments using a Covaris LE220R-plus system (Covaris, USA), followed by end-repair, A-tailing, and adapter ligation. Libraries were amplified by PCR, purified with AMPure XP beads.

Destructive sampling was conducted using skin specimens deposited in the National Museum of Natural History, the Smithsonian Institution (USNM), Field Museum of Natural History (FMNH); American Museum of Natural History (AMNH) and University of Michigan, Museum of Zoology (UMMZ) in the United States. Sterile practices were employed with routine bleaching, instrument sterilization, and changing of PPE to prevent any cross contamination between specimens. Samples of 25 mg of material were placed in sterile cryovials or 1.5mL tubes and transferred to a dedicated Historic DNA processing facility at the Smithsonian Institution’s Museum Support Center and Natural History Museum’s sequencing laboratory where DNA extraction was performed via Qiagen MinElute or Investigator kits following the manufacturer’s protocol. Single-stranded library preparation was performed with the SRSLY Pico Plus Kit (Claret Biosystems®) utilizing half reactions. Libraries had dual indexed iTru style adapters ligated to each sample and amplified for 18 cycles of PCR. Following amplification, samples were visualized on a High-Sensitivity Tape Station (Agilent) and pooled in equimolar ratios for sequencing.

Sequencing was performed on an Illumina NovaSeq X platform to generate 150 bp paired end (PE150) reads. Raw FASTQ reads were processed with fastp v0.24.0 ^76^ to remove adapter contamination and poly-G tails, trimming bases with Phred score ≤ 5. Reads containing > 10 % ambiguous bases or > 50 % low-quality bases were discarded. We supplemented our dataset with 34 available genome re-sequencing data (**Supplementary Table S2**).

### Mitochondrial assembly and phylogenetic analyses

Mitochondrial genomes were assembled from 65 raw sequencing datasets, including data from 31newly sequenced individuals and 34 previously published genomes of various pangolin species. For these datasets, clean reads were processed for *de novo* assembly and annotation in MITOZ v3.6 ^77^, which employed Megahit with k-mer values of 21, 33, and 55. This final set of assembled mitogenomes was then augmented with five additional mitogenomes obtained from GenBank (**Supplementary Table S2**). We aligned mitogenomes using MAFFT v7.5 ^78^, and trimmed poorly aligned regions using Gblocks. We retained 12S, 16S rRNA, and 12 coding genes on the heavy chain for phylogenetic analyses. We also downloaded available mitochondrial 18 *CYTB* and 12 *COI* genes identified as *M. pentadactyla* from GenBank. We used IQ-TREE v2.1.3 to estimate the best-scoring Maximum-likelihood (ML) trees. ModelFinder was used to automatically determine the optimal partitioning scheme and substitution models under the Bayesian Information Criterion (–m MFP+MERGE). Node support was evaluated with 1,000 ultrafast bootstrap replicates.

### Genomic dataset and phylogenetic analyses

We aligned clean data to a *M. pentadactyla* reference genome (GCF_030020395.1) using BWA v0.7.17. The resulting SAM files were converted to coordinate-sorted BAMs and indexed with SAMtools v1.10 ^79^. Duplicates were identified and flagged using MarkDuplicates tool of GATK v.4.6 ^80^. Local realignment around indels was then performed using GATK’s IndelRealigner to reduce false-positive variant calls. Per-sample gVCFs were generated with GATK’s HaplotypeCaller and merged via CombineGVCFs. SNP calling was then executed with GenotypeGVCFs and SelectVariants to produce multisample VCF and isolate high-confidence SNP calls, respectively. We filtered SNP calls using GATK’s VariantFiltration module with thresholds: QUAL < 30.0, QD < 2.0, MQ < 40.0, FS > 60.0, SOR > 3.0, MQRankSum < –12.5, ReadPosRankSum < –8.0, and SB ≥ –1.0. We applied VCFtools v0.1.16 to remove SNPs with minor-allele frequency < 2 % or depth outside the 2.5–97.5 percentile of the depth distribution ^81^. Phylogenetic relationships were inferred using three datasets: high-quality single nucleotide polymorphisms (SNPs) for autosomal and sex chromosomes as well as single-copy orthologous genes. For the SNP-based analyses, autosomal and sex chromosome SNP data were separately extracted from the filtered, high-quality multi-sample VCF. These datasets were converted to PHYLIP format using vcf2phylip v2.8. ML trees were constructed for both datasets using IQ-TREE v2.1.4 as mentioned above. For the ortholog-based analysis, we first generated individual consensus sequences by applying bcftools consensus v1.15. Based on the reference annotation, AGAT was employed to extract the CDS sequences, and translated into protein sequences via EMBOSS transeq v6.6. OrthoFinder v2.5.4 ^82^ initially identified 23,752 single-copy orthologs, which were then subjected to rigorous filtering for GC content, sequence variability, and proportion of missing data. This process yielded a refined set of 7,852 high-quality loci. Each locus was individually aligned and a concatenated ML tree was constructed using IQ-TREE as mentioned above.

### Genetic structuring analyses

To investigate population genetic structure, we utilized linkage-disequilibrium (LD)–pruned autosomal SNPs. LD pruning was performed in PLINK v1.90b6.21 using the –indep-pairwise 50 5 0.2 parameters ^83^. We then performed PCA on mean-imputed genotypes in PLINK to extract the top ten principal components. To assess the statistical significance of the principal components, we applied the Tracy-Widom test implemented in EIGENSOFT v8.0.0 ^84^. We applied PCA to all *Manis* samples and to *M. pentadactyla* sensu lato (which encompasses individuals later identified as *M. aurita*). To infer individual ancestry proportions, we ran ADMIXTURE v1.3.0 on the *M. pentadactyla* s.l. samples. We evaluated models for K (number of ancestral populations) from 1 to 10, and selected the optimal K to minimize cross-validation error ^85^.

### Molecular dating and population genetic analyses

Divergence times were estimated using *BEAST3 model ^86^ as implemented in BEAST v2.7.7 ^87^. First, we identified the 2,000 most genetically divergent loci from an initial pool of 7,852 filtered orthologs. These loci were then assessed for their conformity to a strict molecular clock using SortaDate v2.0 ^88^. The 200 orthologs that best fit the strict clock model were subsequently used for the divergence time estimation. The Bayesian inference employed a Birth–Death tree prior, a strict molecular clock, and the HKY+Γ substitution model. Given the scarcity of unambiguous crown-group fossils for extant pangolin, we calibrated the time tree using a combination of secondary and fossil-based constraints. First, we applied two secondary calibrations derived from a genome-scale pangolin dating analysis Gu, et al. ^10^, which estimated (i) the basal split between African and Asian pangolins at 33.19 Ma (30.34–39.80 Ma) and (ii) the divergence of crown Asian pangolins (genus *Manis*) at 16.85 Ma (12.92–22.36 Ma). We incorporated a fossil-based minimum constraint within African pangolins (*Smutsia gigantea* and *Phataginus tricuspis*) using the earliest confirmed fossil representative of the giant pangolin lineage from the Varswater Formation at Langebaanweg, South Africa, dated to the early Pliocene (∼5 Ma) ^89^. The Markov chain Monte Carlo (MCMC) analysis was run for 300 million generations. Convergence and parameter mixing were assessed in Tracer v1.7.2 ^90^, ensuring all key parameters achieved effective sample sizes (ESS) > 200.

Historical population dynamics were reconstructed using the Pairwise Sequentially Markovian Coalescent (PSMC) model ^91^. According to the estimated divergence time between *M. pentadactyla* and *M. aurita* (see Results), we estimated the mutation rate per generation assuming an average generation time of 7 years ^92^. The analysis was run with the parameters -N25 -t15 -r5 - p “4+25*2+4+6”, using the Manis *pentadactyla* genome ^13^ as the reference for mapping. To assess the variability in demographic inferences, 100 bootstrap replicates were performed.

To complement the long-term demographic reconstructions from PSMC, we inferred recent effective population size (Ne) dynamics over the last ∼600 generations for *M. aurita* and *M. pentadactyla* using the linkage-disequilibrium method implemented in GONE ^93^. We used the same autosomal whole-genome SNP dataset filtered for high genotype quality and site callability. To evaluate robustness, we generated 100 replicate datasets by random subsampling, limiting each chromosome to a maximum of 100,000 SNPs to standardize marker density among chromosomes. Each replicate was analyzed under identical GONE settings (cMMb = 1, maxNSP = 50,000, hc = 0.05), and replicate results were summarized to quantify uncertainty in the inferred N_e_ trajectories.

Levels of inbreeding were quantified by calculating the average inbreeding coefficient (F_ROH_) for each individual. Long runs of homozygosity (ROH) were identified using PLINK v2.0 ^83^ with the parameters --homozyg --homozyg-window-snp 20 --homozyg-kb 100. Individual F_ROH_ values were then calculated as the proportion of the autosomal genome covered by ROH, relative to the total autosomal SNP coverage, following McQuillan, et al. ^94^. Because DNA from museum specimens is often degraded and highly fragmented, long runs of homozygosity (ROHs) may be under-detected, potentially biasing F_ROH_ estimates downward. We therefore restricted ROH and F_ROH_ analyses to contemporary samples.

### Geneflow analyses

To infer the direction and magnitude of historical gene flow among *Manis* species, we analyzed a high-quality SNP dataset from 52 samples representing the four *Manis* species using Dsuite v0.5 ^95^ and TreeMix v1.13 (Pickrell & Pritchard, 2012). We first filtered the SNP dataset to retain sites with ≤20% missing data using VCFtools (--max-missing 0.8). To quantify admixture, we used the Dtrios program in Dsuite to calculate D-statistics and f4-ratio statistics for all relevant population trios. *P. tricuspis* and *S. gigantea* were used as outgroups. Results were visualized with the dtools.py script. To model population splits and migration events, the filtered SNP data was first pruned for linkage disequilibrium (PLINK; --indep-pairwise 50 5 0.6), and the output was converted to TreeMix format using plink2treemix.py. We then ran TreeMix, exploring models with 0 to 5 migration edges (five replicates each). The optimal model was selected by evaluating the explained variance and residual plot fits. The resulting population graph was visualized using the R script plotting_funcs.R provided with TreeMix.

We used FastSimCoal v2.8 ^96^ to test potential gene flow between *M. pentadactyla* (MP) and *M. aurita* (MA). First, we used VCF file through easySFS to generate folded observed joint site frequency spectrum (JSFS). We constructed four demographic models to test gene flow scenarios: (1) early gene flow (range from 120,000 to 130,000 generations); (2) recent gene flow (range from 1,000 to 20,000 generations); (3) continuous gene flow; and (4) no gene flow. For each tested demographic model, we performed 100 independent runs using 100,000 simulations, 40 expectation–maximization cycles, and a broad search range for parameters to determine the run with the best parameter estimates and maximum likelihood. Model comparisons were performed using the Akaike Information Criterion (AIC) to identify the model that best explains the observed data.

### Genomic Island of Differentiation

To identify genomic regions under divergent selection, we scanned the genomes of *M. pentadactyla* and *M. aurita*. We used Pixy v2.0 to calculate nucleotide diversity (π), fixation index (F_ST_), and absolute genetic differentiation (D_XY_) across 20 kb windows ^97^. Candidate genomic islands of divergence were defined as outlier windows concurrently in the top 5th percentile for both F_ST_ and D_XY_. We extracted all genes within these outlier regions. Genes annotated only with LOC identifiers were annotated based on their high-confidence human orthologs, identified via BLASTP ^98^. This gene set was then tested for Gene Ontology (GO) term enrichment using the R package clusterProfiler ^99^.

### Morphometric analyses

We accessed specimens deposited in FMNH, USNM, UMMZ, British Museum of Natural History, UK (BMNH), Natural History Museum, Tribhuvan University (NHM-TU) Institute of Zoology, Guangdong Academy of Sciences, China (IOZ-GDAS), and Guangdong Wildlife Monitoring and Rescue Center, China (GWMRC) (**Supplementary Table S1**). Fourteen external measurements were taken including full body length (abdominal) (FBLA), head body length (abdominal) (BLA), tail length (abdominal) (TLA), full body length (back) (FBLB), head body length (back) (HBL), tail length (back) (TLB), external pinna length (EL), external pinna thickness (ET), middle claw length of the back foot (MCLBF), middle claw length of the forefoot (MCLFF), the distance of the first scale from the tip of the nose (DFSTN), the maximum number of scales in the back of the body (MNS), the single-sided number of tail edge scales (SNTE), length of hind foot (LHF), using soft tape rulers (accurate to 0.1 mm) for flexible dimensions and MNT-150 standard-type zinc alloy digital caliper (Model: MNT952101; DEGUQMNT, accuracy to 0.01 mm) for more precise linear features (**Supplementary Figure S3A**). This data includes 26 skin specimens for *M. pentadactyla* (n=8), *M. javanica* (n=12), *M. aurita* (n=6) and also includes the lectotype of *M. aurita* (**Supplementary Data S1**).

Twenty cranial measurements were conducted with MNT-150 standard-type zinc alloy digital caliper (accuracy to 0.01 mm), following ^30,100–104^ including total skull length (TSL), nasal to foramen magnum base length (NFBL), length from premaxilla to palatine (PP), from Intersection between maxilla and palatine sutures to most posterior point of the palatine sutures(IMPS), tymapanic bulla (left) (TB), skull width (SW), skull height (SH), maximum width of nasal bone (WN), maximum length of nasal bone (LN), dorsal gap between maxillary (dorsal) (GM), maximum width of gap between alveolar sulci (MWG), left maximal length of orbit left, (MO), distance between tympanic bulla (DB), length of suture between parietal bone (DTEM), length of suture between frontal bone (DLOB), Length of the zygomatic process between maxilla and squamous bone (left) (ZMS), length of foremen magnun occipitals (FMO), mandible length (left) (ML), the distance between condylar process (DCP), length of palatal process of premaxilla(LPP) (**Supplementary Figure S3B**). This data includes 44 skulls for *M. pentadactyla* (n=25), *M. javanica* (n=7), *Manis crassicaudata* (n=2)*, M. culionensis* (n=3) and *M. aurita* (n=7; **Supplementary Data S2**). Sex was not considered in the analysis due to the absence of marked sexual dimorphism and the lack of information for some specimens. Specimen age was assessed based on body size, skull morphology, and available collection records. Juvenile and subadult individuals were excluded in the analysis. To ensure completeness of the data, we replaced missing values with the mean value calculated across specimens of the same species following <u>Arbour and</u> Brown ^105^. We used the function prcomp () of stat to conduct PCA and Morpho to perform CVA on log₁₀-transformed variables. Finally, we conducted Levene’s test for homogeneity of variances using the leveneTest () function and then performed Student’s t-tests with t.test () to compare group means across taxa.

We downloaded 84 micro-CT scan data from Ferreira-Cardoso, et al. ^106^, and scan 13 specimens, obtaining high-resolution micro-CT scans of 97 skulls for *M. pentadactyla* (n = 42), *M. javanica* (n = 38), *M. crassicaudata* (n = 12), *M. culionensis* (n = 3), and *aurita* (n = 2; **Supplementary Table S2**). Raw DICOM stacks were imported into Materialise Mimics v19.0 for segmentation and construction of 3D cranial meshes. We placed 75 landmarks per mesh following Ferreira-Cardoso (2020; **Supplementary Figure S5**). After Procrustes alignment, we conduct PCA and CVA using MorphoJ v1.08 ^107^. Landmark selection was based on prior studies addressing mammalian taxa ^108,109^

### Scale coding

We generated scale-encoded heatmaps from 12 intact skins (nine *M. pentadactyla,* three *M. aurita*) to visualize variation following Algewatta, et al. ^110^; Guo, et al. ^111^ and Martin, et al. ^112^. In brief, we recorded the x-y coordinates of every scale in Excel, exported standardized images to Photoshop v25, and overlaid all specimens within each species to generate species-specific composite maps (**Supplementary Data S3 and S4**).

## DATA ACCESSIBILITY

All sequencing data were deposited in the National Center for Biotechnology Information (NCBI) database (BioProjectID <u>PRJNA1332899</u>). Sample metadata, including individual identifiers and accession numbers, are summarized in **Supplementary Table S2**. The analysis scripts and source data for the main figures are available on Dryad repository with the identifier Dryad http://datadryad.org/share/tUnpoAtBaYbC-LxLmopIeUXwwngHX96olu7F6k_uTBk ^113^

## SUPPLEMENTARY DATA

Supplementary data to this article can be found online.

## COMPETING INTERESTS

The authors declare that they have no competing interests

## AUTHOR CONTRIBUTIONS

**NPK and KH**: Conceptualization, Methodology, Investigation, Resources, Visualization, Validation, Supervision, Funding acquisition, Project administration, Writing – review & editing. All authors read and approved the final version of the manuscript. **ZZ, GZ, XH, AF, and YH**: Methodology, Investigation, Formal analysis, Visualization, Validation, Writing – original draft, Writing – review & editing. **ZY, XW, JL, DM, SB, LH, WY, and LL** Methodology, Investigation, Formal analysis, Visualization, Validation. **MTRH, CG, AH, and MFCF,** Formal analysis, Validation, Writing – original draft, Writing – review & editing.

## ACKNOWLEDGMENTS

We thank the following institutions for providing access to specimens for morphological and genetic analyses: Natural History Museum (NHM), UK, the National Museum of Natural History, Smithsonian Institution (USNM); Field Museum of Natural History (FMNH); American Museum of Natural History (AMNH); University of Michigan Museum of Zoology (UMMZ); the Institute of Zoology, Guangdong Academy of Sciences (IOZ-GDAS); and the Guangdong Wildlife Monitoring and Rescue Center (GWMRC), China. We are grateful to the Natural History Museum, Tribhuvan University, and the Department of National Parks and Wildlife Conservation, Nepal, for providing research permits and institutional support. We sincerely thank the Ministry of Education, Science and Technology (MoEST), Nepal, and the Nepal Academy of Science and Technology (NAST) for their support. We also thank Dr Dmitry Telnov and Roberto Portela Miguez, curators at NHM UK for their assistance.

NPK’s effort to this study was partially supported by MoEST, Nepal, 2019, an ANSO-CAS visiting Scientist fellowship at Kunming Institute of Zoology, Chinese Academy of Sciences, 2025 and South China Biodiversity Research Center, School of Life Sciences, Guangzhou University, Guangzhou, China, 2023. This study is supported by the ADCS Seed Grants program of the National Museum of Natural History, Smithsonian Institution (M.T.R.H. and A.H.), Guangdong Provincial Natural Science Foundation for Distinguished Young Scholars (2022B1515020033, K.H.) and the National Natural Science Foundation of China (32170452, K.H.).

## Supplementary figures

Supplementary Figure S1. Maximum likelihood phylogeny of pangolins based on sex-chromosome SNPs. The tree was constructed using the Maximum Likelihood (ML) method in IQ-TREE from an alignment of single-nucleotide polymorphisms (SNPs) derived from sex chromosomes. Numbers at the nodes represent bootstrap support values from 1000 replicates. The scale bar indicates the number of substitutions per site. The analysis shows that *Manis aurita* and *Manis pentadactyla* s.s. each form a distinct, reciprocally monophyletic clade with maximum support.

Supplementary Figure S2. Maximum likelihood phylogeny based on single-copy orthologous genes. The tree was inferred using the Maximum Likelihood method in IQ-TREE from a concatenated dataset of 7,852 single-copy orthologous genes. Numbers at nodes represent bootstrap support values from 1,000 replicates (only maximum support values of 100 are shown). The scale bar indicates the number of substitutions per site. Clades are colored according to the species legend.

Supplementary Figure S3. Illustration of morphometric measurements taken from the pangolin body and cranium. (A) External body measurements. (B) Cranial and mandibular measurements shown in ventral and dorsal views. The pangolin image is adopted from Ognimdo, 2002. Abbreviations for external body measurements (A): FBLA, full body length (abdominal); BLA, head body length (abdominal); TLA, tail length (abdominal); FBLB, full body length (back); HBL, head body length (back); TLB, tail length (back); EL, external pinna length; ET, external pinna thickness; MCLBF, middle claw length of the back foot; MCLFF, middle claw length of the forefoot; DFSTN, distance of the first scale from the tip of the nose; MNS, maximum number of scales on the back of the body; SNTE, single-sided number of tail edge scales; LHF, length of hind foot. Abbreviations for craniodental measurements (B): DB, Distance between tympanic bulla; DBCP, Distance between condylar processes; DLOB, Length of the suture between frontal bones; DTEM, Length of the suture between parietal bones; FMO, Length of the foramen magnum; GM, Gap between maxillary bones (dorsal); SH, Skull height; IMPP, Length from the intersection of maxilla-palatine sutures to the most posterior point of the palatine; LN, Maximum length of the nasal bone; LPP, Length of the palatal process of the premaxilla; ML, Mandible length; MO, Maximal length of the orbit; MWG, Maximum width of Gap between alveolar sulci; NFBL, Length from the nasal to the foramen magnum base; PP, Length from the premaxilla to the palatine; TB, Tympanic bulla length; TSL, Total skull length; WN, Maximum width of the nasal bone; SW, skull width; ZMS, Length of the zygomatic process between the maxilla and squamosal bones (left).

Supplementary Figure S4. Pairwise plots of Principal Component Analyses (PCA) for three morphological datasets. In all plots, points are colored by species, and shaded polygons represent the convex hull for each species group. Values in parentheses on the axes indicate the percentage of variance explained by each principal component. (A, B) Plots of PC1 vs. PC3 and PC2 vs. PC3 from the PCA based on 20 linear cranial measurements. (C, D) Plots of PC1 vs. PC2 and PC1 vs. PC3 from the PCA of cranial shape variation derived from 75 3D geometric morphometric landmarks. (E, F) Plots of PC1 vs. PC3 and PC2 vs. PC3 from the PCA based on 14 external body measurements. Note the inclusion of the *M. aurita* holotype (BMNH 43.1.12.85), which is labeled in these plots.

Supplementary Figure S5. Location of the 75 three-dimensional landmarks used for the geometric morphometric analysis of the pangolin cranium. The landmarks are shown in (A) lateral, (B) ventral, and (C) dorsal views. This landmarking scheme was adapted from Ferreira-Cardoso et al. (2020). A complete list and description of these landmarks can be found in Table S6.

Supplementary Figure S6. Univariate comparisons of all cranial and external measurements between *M. pentadactyla s.s.* and *M. aurita*. (A) Boxplots of cranial measurements. (B) Boxplots of external body measurements. In all plots, boxes depict the interquartile range (IQR) with the median indicated by the central horizontal line; whiskers extend to 1.5 times the IQR, and outliers are shown as individual points. The y-axis represents the measurement values. For cranial measurements (A), all units are in millimeters (mm). For external measurements (B), units are in centimeters (cm), except for scale counts which are unitless. Statistical significance from Student’s t-tests is denoted as: ns, not significant; * *P* < 0.05; ** *P* < 0.01; *** *P* < 0.001.

Supplementary Figure S7. Canonical Variate Analyses (CVA) of *Manis* specimens based on different morphological datasets. In all plots, points are colored by species. Values in parentheses on the axes indicate the percentage of between-group variance explained by each canonical variate. (A-C) Pairwise plots of the first three canonical variates from the CVA based on 20 linear cranial measurements. (D-F) Pairwise plots of the first three canonical variates from the CVA based on cranial shape variation derived from 75 3D geometric morphometric landmarks. (G) Plot of the first two canonical variates from the CVA based on 14 external body measurements. Note that all three analyses show a clear and highly significant separation between *M. aurita* and *M. pentadactyla s.s., and that the M. aurita holotype (BMNH 43.1.12.85) is correctly classified with other M. aurita specimens in panel G*.

Supplementary Figure S8. Heatmap analysis comparing dorsal scale patterns between *M. aurita* and *M. pentadactyla s.s.* The heatmaps were generated by superimposing scale-encoded images from 12 intact body skins (*n* = 3 for *M. aurita*; *n* = 9 for *M. pentadactyla s.s.*). In each map, the color intensity corresponds to the frequency of a scale being present at that coordinate across all specimens of a species, with darker colors indicating a higher frequency. (A) Composite heatmap for *M. aurita*. (B) Composite heatmap for *M. p. pentadactyla s.s.* (C) Comparative heatmap overlaying the patterns of both species (purple for *M. aurita*, red for *M. pentadactyla s.s.*). The analysis revealed no significant differences in the spatial distribution or pattern of scales between the two taxa.

Supplementary Figure S9. Comparative GONE-inferred effective population size (Ne) histories of *Manis auritus* and *Manis pentadactyla*, showing 95% confidence intervals

Supplementary Figure S10: Genomic landscapes of diversity and divergence between *M. aurita* and *M. pentadactyla s.s.* All statistics were calculated in non-overlapping 20 kb windows. In all panels, each point represents a single window, and the x-axis shows the position across the 19 chromosomes. (A) Nucleotide diversity (π) within *M. pentadactyla s.s.* (B) Nucleotide diversity (π) within *M. aurita*. (C) Population differentiation, measured by the fixation index (*F*_ST_), between the two species. (D) Absolute genetic divergence (*d*_XY_) between the two species.

Supplementary Figure S11. Diagnostic characteristics of the external morphology and cranium of *Manis aurita*, with comparative live photographs of four *Manis* species. (A) Lateral view of the head of specimen NHMTU-NP06, highlighting features 1 and 2. (B) Close-up of dorsal scales from the holotype BMNH 43.1.12.85 (left) and specimen NHMTU-NP07 (right), highlighting feature 3. (C) Ventral view of the cranium of NHMTU-NP04, highlighting features 4, 5, 6, and 8. (D) Lateral view of the cranium of NHMTU-NP04, highlighting feature 7. The numbered diagnostic features are as follows: 1) The external pinna is flap-shaped and distinctly smaller than that of *M. pentadactyla*. 2) On the snout, the first row of head scales is set back from the nasal tip, exposing a patch of bare skin. 3) The medial rows of dorsal scales are broadly trapezoidal with straight posterior margins; lateral scales near the abdomen are broad and shield-shaped. 4) The palatal process of the premaxilla extends posteriorly but measures less than half the premaxillary midline length. 5) The incisive foramen is oval-shaped and enclosed entirely within the premaxilla, not reaching the posterior border nor contacting the maxilla. 6) The maxilla is laterally inflated at its anterior end, producing a robust rostral profile with nearly parallel lateral margins. 7) The zygomatic arch is complete or nearly complete, with the maxillary and squamosal zygomatic processes contacting or separated by a narrow gap. 8)The postglenoid foramen opens laterally on the squamosal zygomatic process. 9) The jugular and hypoglossal foramina open within a single depressed fossa on the ventral surface of the cranium. **(E–F)** Live head photographs of two *Manis* species highlighting diagnostic features of the external pinna: **(E)** *Manis aurita* from Kavre, Nepal. Photograph by Prabin Shrestha. **(F)** *Manis pentadactyla* from Guangzhou, China. Photograph by Hua Yan. **(G–J)** Live habitus photographs of four extant *Manis* species for comparison: **(G)** *Manis aurita* from Kavre, Nepal. Photograph by Prabin Shrestha. **(H)** *Manis pentadactyla* from Guangzhou, China. Photograph by Hua Yan. **(I)** *Manis javanica* from Guangzhou, China. Photograph by Hua Yan. **(J)** *Manis crassicaudata* from Kailali, Nepal. Photograph by prabin Shrestha.

Supplementary Figure S12. Cranial and external morphology among five *Manis* species. (A) Cranial morphology shown in ventral, lateral, and dorsal views. The numbered features shown are: 1) The incisive foramen of the premaxilla and its relationship to the ventral projection of the maxilla. 2) Extent of posterior extension of the palatal process of the premaxilla. 3) Degree of ventral expansion of the maxilla. 4) Whether the jugular and hypoglossal foramina are recessed. 5) Orientation of opening of the postglenoid foramen. (B) Dorsal skull scales.

Supplementary Figure S13. Maximum likelihood phylogeny of *Manis* species, incorporating previously published sequences. The tree was inferred using the Maximum Likelihood (ML) method in IQ-TREE from a combined dataset of complete mitochondrial genomes and partial CYTB and COI gene sequences. Numbers at the nodes represent bootstrap support values from 1,000 replicates. The scale bar indicates the number of substitutions per site. Major clades are colored according to the species legend. Individuals with tip labels shown in red are sequences retrieved from Xie et al. (2021), and this analysis confirms their phylogenetic placement within the *M. aurita* and *M. pentadactyla s.s.* clades.

## Supplementary tables

Table S1. List of Specimens Used in All Morphological Analyses.

Table S2. Detailed Information for All Specimens Included in The Genetic/Genomic Analyses, Including Voucher Numbers, Locality, Data Type, and Source.

Table S3. Summary Statistics and Quality Metrics for The Whole-Genome Resequencing Data.

Table S4. Results of Principal Component Analyses (PCA) and Associated Significance Tests for Morphological and Genomic Data.

Table S5. Cross-Validation Errors for Different K Values in The ADMIXTURE Analysis.

Table S6. Anatomical Definitions of the 75 Three-Dimensional Landmarks Used in The Geometric Morphometric analysis.

Table S7. Detailed Results of The Principal Component Analysis (PCA) Based on 20 linear Cranial Measurements.

Table S8. Detailed Results of The Principal Component Analysis (PCA) Based on 3D Geometric Morphometric Data.

Table S9. Detailed Results of The Principal Component Analysis (PCA) Based on 14 External Body Measurements.

Table S10. Results of Independent-Samples T-Tests for Cranial Measurements Between *M. pentadactyla* s.s. and *M. aurita*.

Table S11. Results of Independent-Samples T-Tests for External Body Measurements Between *M. pentadactyla* s.s. and *M. aurita*.

Table S12. Detailed Results of The Canonical Variate Analysis (CVA) Based on 20 linear Cranial Measurements.

Table S13. Detailed Results of The Canonical Variate Analysis (CVA) Based on 14 External Body Measurements.

Table S14. Detailed Results of The Canonical Variate Analysis (CVA) Based on 3D Geometric Morphometric Data.

Table S15. Information on T-Test for Scale Counts

Table S16. Results of D-Suite Analyses Testing for Interspecific Gene Flow.

Table S17. Likelihood Comparison of Alternative Demographic Models of Gene Flow between M. aurita and M. pentadactyla s.s.

Table S18. Morphological Character Matrix of Asian Pangolins (Genus *Manis*).

## Supplementary Data

Data S1. External Body Measurements of Pangolins

Data S2. Length of suture between parietal bone (DTEM)

Data S3. Scale Positioning Data of pangolins

Data S4. Scale Counts in Different Body Regions of Pangolins

## Notes

http://datadryad.org/share/tUnpoAtBaYbC-LxLmopIeUXwwngHX96olu7F6k_uTBk

### Competing Interest Statement

The authors have declared no competing interest.

### Summary of Updates

Revised manuscript has updated figures, tables. We have added new WGS data from lectotype.

